# Polygenic prediction of breast cancer: comparison of genetic predictors and implications for screening

**DOI:** 10.1101/448597

**Authors:** Kristi Läll, Maarja Lepamets, Marili Palover, Tõnu Esko, Andres Metspalu, Neeme Tõnisson, Peeter Padrik, Reedik Mägi, Krista Fischer

**Author notes:** Correspondence:* Kristi Läll, Address: Riia 23b, Tartu, 51010, Estonia.

## Abstract

**Background:** Published genetic risk scores for breast cancer (BC) so far have been based on a relatively small number of markers and are not necessarily using the full potential of large-scale Genome-Wide Association Studies. This study aims to identify an efficient polygenic predictor for BC based on best available evidence and to assess its potential for personalized risk prediction and screening strategies.

**Methods:** Four different genetic risk scores (two already published and two newly developed) and their combinations (metaGRS) are compared in the subsets of two population-based biobank cohorts: the UK Biobank (UKBB, 3157 BC cases, 43,827 controls) and Estonian Biobank (EstBB, 317 prevalent and 308 incident BC cases in 32,557 women). In addition, correlations between different genetic risk scores and their associations with BC risk factors are studied in both cohorts.

**Results:** The metaGRS that combines two genetic risk scores (metaGRS_2_ - based on 75 and 898 Single Nucleotide Polymorphisms, respectively) has the strongest association with prevalent BC status in both cohorts. One standard deviation difference in the metaGRS2 corresponds to an Odds Ratio = 1.6 (95% CI 1.54 to 1.66, *p* = 9.7*10^-135^) in the UK Biobank and accounting for family history marginally attenuates the effect (Odds Ratio = 1.58, 95% CI 1.53 to 1.64, p = 9.1*10^-129^). In the EstBB cohort, the hazard ratio of incident BC for the women in the top 5% of the metaGRS_2_ compared to women in the lowest 50% is 4.2 (95% CI 2.8 to 6.2, p = 8.1*10^-13^). The different GRSs are only moderately correlated with each other and are associated with different known predictors of BC. The classification of genetic risk for the same individual may vary considerably depending on the chosen GRS.

**Conclusions:** We have shown that metaGRS_2_ that combines on the effects of more than 900 SNPs provides best predictive ability for breast cancer in two different population-based cohorts. The strength of the effect of metaGRS2 indicates that the GRS could potentially be used to develop more efficient strategies for breast cancer screening for genotyped women.

## Background

Breast cancer (BC) is the most frequent cancer among women in the world, being also the second leading cause of cancer death in women in more developed regions after lung cancer^1^. As early diagnosis for BC could lead to successful treatment and good prognosis for recovery, it is important to develop efficient risk prediction algorithms that aid to identify high-risk individuals. Although many countries have implemented mammography screening programs, they mostly apply to all women in certain age categories without any additional stratification by other risk factors. However, the benefits of such screening programs are often debated. Existing tools to assess BC risk^2-4^ are often not systematically used in screening due to insufficient up-to-date risk factor’s information. Also, they only capture the heritable component either in the form of family history or using the information on rare genetic variants (BRCA1/2).

It has been estimated in twin studies that the heritability of breast cancer ranges from 20 to 30%^5^. However, only 5%–10% of BC cases have a strong inherited component identified in a form of rare genetic variants^6^, indicating that in addition there should be a considerable polygenic component in the disease liability. This is also supported by the results of large genome-wide association studies (GWAS) – more than 100 genomic loci have been identified as being associated with BC in Europeans^7^.

Based on the GWAS results, several efficient polygenic risk scores (GRS) have been developed for common complex diseases that in many cases can be used to improve the existing risk prediction algorithms^8-11^. It is natural to expect that a similar GRS for BC may aid risk prediction in clinical practice.

So far, several studies have combined the SNPs with established genome-wide significance in a GRS for BC. Sieh *et al*^12^ used 86 SNPs and Mavaddat *et al*^13^ 77 SNPs to calculate a GRS, both showing a strong effect of the score in predicting future BC cases. Few studies have also demonstrated the incremental value of adding GRS to proposed BC prediction algorithms^14,15^. Although several different GRSs have been proposed for BC risk prediction, no head-to-head comparison of the scores has been found in the literature. It has also not been assessed, whether the number of SNPs in the GRS could be increased. The latter was also problematic due to unavailability of summary statistics from large-scale GWASs.

In 2017, the large scale GWAS by Michailidou *et al*^7^ released summary statistics for around 11.8 million genetic variants. Almost at the same time, UK Biobank released their GWAS results for BC for ∼10.8 million SNPs. As evidence from studies on other common complex diseases indicates that predictive ability of a GRS can improve by adding the effects of a large number of independent SNPs in addition to the ones with established genome-wide significance, we intend to explore this approach using both summary files.

## Methods

### Study cohorts

In the present analysis, the data of 32,557 female participants of the Estonian Biobank (EstBB)^16^ has been used, with 317 prevalent and 308 incident cases of BC. Incident disease data was obtained from linkages with the Estonian Health Insurance Fund, Estonian Causes of Death Registry and Estonian Cancer Registry (latest update in December 2015).

We have also analyzed the data of 46,984 women (incl 3,157 BC cases) of European ancestry from the UK Biobank^17^ who passed the main quality control and were not included in the UKBB breast cancer GWAS^18^.

More details about cohorts can be found in the Additional File 2 and overview of the characteristics of the cohorts is given in the Additional File 1, Table S1.

### Statistical Methods

#### General concept of Genetic Risk Scores (GRS)

The general definition of a GRS is based on the assumption that the polygenic component of the trait (e.g. disease risk) can be approximated by a linear combination of *k* independent SNPs

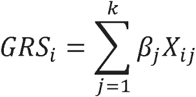

where *β_j_* is the weight of each SNP and *X_ij_* represents the number of risk alleles for *j* – *th* SNP(*j* = 1,…,*k*) for the *i* – *th* individual, (*i* = 1,…,*n*) Typically the estimated (logistic) regression coefficients from a large-scale GWAS meta-analysis are used as weights *β_j_*.

Published versions of GRS can be divided to two main categories. We call a GRS *multigenic,* if the number of SNPs (*k*) is relatively small, containing only the SNPs with established genome-wide significance from a GWAS. A *polygenic* GRS may contain a large number of SNPs (often *k* > 1000) and is either based on all available independent SNPs (with pairwise correlation not exceeding a pre-defined threshold) or the ones that satisfy some p-value threshold (often ≥ 0.05).

In the present paper, we will compute two multigenic and two polygenic GRSs, whereas the polygenic GRSs are developed using the PRSice software^19^.

#### Computation of multigenic and polygenic GRSs and analysis of their association with prevalent breast cancer

First we calculate two previously published multigenic GRSs for the EstBB data – both scores contain only those SNPs from the originally published versions that are available with acceptable imputation accuracy in the EstBB.

1. The score denoted by **GRS**_70_, based on Sieh *et al*^12^ (70 SNPs out of 86 were available).
2. The score GRS_75_, based on the 75 SNPs of the 77-SNP score by Mavaddat *et al*^13^. Next, two polygenic GRSs were developed. For both GRSs, first a set of SNPs was created so that: a) GWAS summary statistics are available for the entire set; b) the SNPs are genotyped or imputed with an acceptable quality in the EstBB; c) the SNPs are independent – the pairwise correlation does not exceed a pre-specified threshold (details on SNP selection provided in the Additional File 2). For the final selection of the p-value threshold for the SNPs to be included in the GRS, age-adjusted logistic regression model comparing 317 prevalent BC cases and 2000 randomly chosen controls in the EstBB cohort was used and the score with the smallest p-value for the GRS-phenotype association was selected. The resulting polygenic scores are:
3. The score **GRS**_ONCO_, based on the summary statistics of the Breast Cancer Association Consortium meta-analysis of BC with 122,977 cases and 105,974 controls^7^.
4. The score **GRS**_UK_, based on the summary statistics of the GWAS conducted on the UK Biobank data (comparing 7,480 BC cases and 329,679 controls including both men and women^18^). The reported linear regression coefficients were transformed into corresponding log odds ratios, following the rules described by Lloyd-Jones *et al^20^*, before using them as weights in the GRS.
5. Thereafter, Pearson coefficients of correlation between different GRSs were calculated. The GRSs were combined into three different versions of metaGRS, following the ideas by Inouye *et al*^21^: **metaGRS**_4_ as the weighted average of all four GRSs, **metaGRS**_3_ as the weighted average of three GRSs with the strongest association with incident BC and finally **metaGRS**_2_ based on top two predicting GRSs. As weights to construct metaGRS, log(odds ratios) of GRSs from training set from logistic regression model were used.

Finally, the UK biobank data was used to address the attenuation of GRS’ effect while accounting for family history of BC and to study associations between BC risk factors and GRSs. While modelling in UK biobank, age at recruitment and 15 principal components are included in the model.

#### Analysis of the GRS effects on incident BC

All 7 GRSs were evaluated in the analysis of incident BC in 30240 women from the EstBB cohort who did not have an existing BC diagnosis at recruitment and were not included in the case-control set used to select the best polygenic GRSs. Cox proportional hazard models were used to estimate the crude and adjusted Hazard Ratios (HR) corresponding to one standard deviation (SD) of the GRS. To properly account for left-truncation in the data, age of the participant was used as timescale in the analyses. To assess the incremental value of GRSs when added to other known risk factors, the models were additionally adjusted for the absolute risk estimates from the NCI Breast Cancer assessment tool^2,22^, based on age, race, age at menarche and age at first live birth of the participant. Other possible risk factors such as number of biopsies were set as unknown. Harrell’s c-statistic to characterize the discriminative ability of each GRS and their incremental value compared to NCI’s Breast Cancer assessment tool absolute risk estimates alone were computed. Hazard ratios for GRS top quintile and top 5% percentile compared to average and low GRS categories were reported. Cumulative incidence estimates were computed with Aalen-Johansen estimator to account for competing risk.

Finally, associations between GRSs and variables related to female’s reproductive health and BC risk factors are explored using linear, logistic or Cox regression models depending on the type of dependent variable in both EstBB and UKBB cohorts (more details in the Additional File 2).

## Results

### GRSs association with prevalent breast cancer

Both GRS70 and GRS75 were significantly associated with prevalent BC status in the case-control subset of the EstBB cohort, with corresponding Odds Ratio(OR) estimates per one SD of the GRS being 1.27 (95% CI 1.13 to 1.45, p = 1.4*10^-4^) and 1.38 (95% CI 1.22 to 1.57, p = 5.3*10^-7^), respectively. Of all polygenic GRSs, the strongest association was observed for GRS_ONCO_ with p-value threshold p <5* 10^-4^ for SNP inclusion (898 SNPs). This resulted in OR = 1.44 (95% CI 1.27 to 1.64, p = 1*10^-8^) per one SD of the GRS. The best version of GRS_UK_ included 137 SNPs that satisfied inclusion threshold p<5*10^-5^ and resulted in OR = 1.34 (95% CI 1.18 to 1.52, p = 5.5*10^-6^). Similar effect sizes for all four GRSs were observed in the UKBB cohort (Table S2). Detailed results on GRS-outcome associations in EstBB with different p-value thresholds for SNP inclusion can be seen in Additional File 2, Figure S1.

### Association of incident breast cancer and GRSs

Out of four studied GRSs, GRS_UK_ has the weakest and GRS75 the strongest association with incident BC (Table 1) in the EstBB, both in terms of the p-value as well as the Harrell’s c-statistic. All metaGRSs have stronger association with incident BC than original scores alone. However, when GRS_ONCO_ and GRS_75_ are already combined into metaGRS_2_, no additional gain is seen from adding GRS_UK_ and/or GRS_70_ to the score. Therefore, we chose metaGRS_2_ for further assessment of its properties. While a predictive model capturing the effect of the NCI risk estimates resulted in the Harrell’s c-statistic of 0.677, it was increased to 0.715 (by 3.8%) when also metaGRS_2_ was added to the model.

**Table 1.**
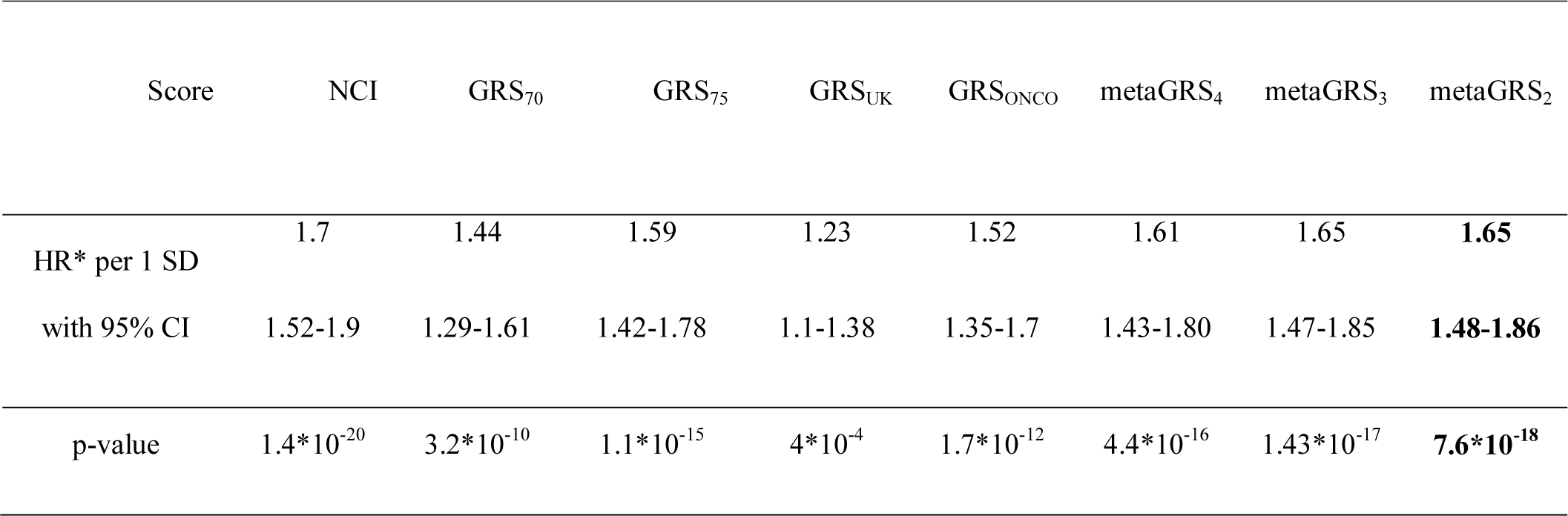
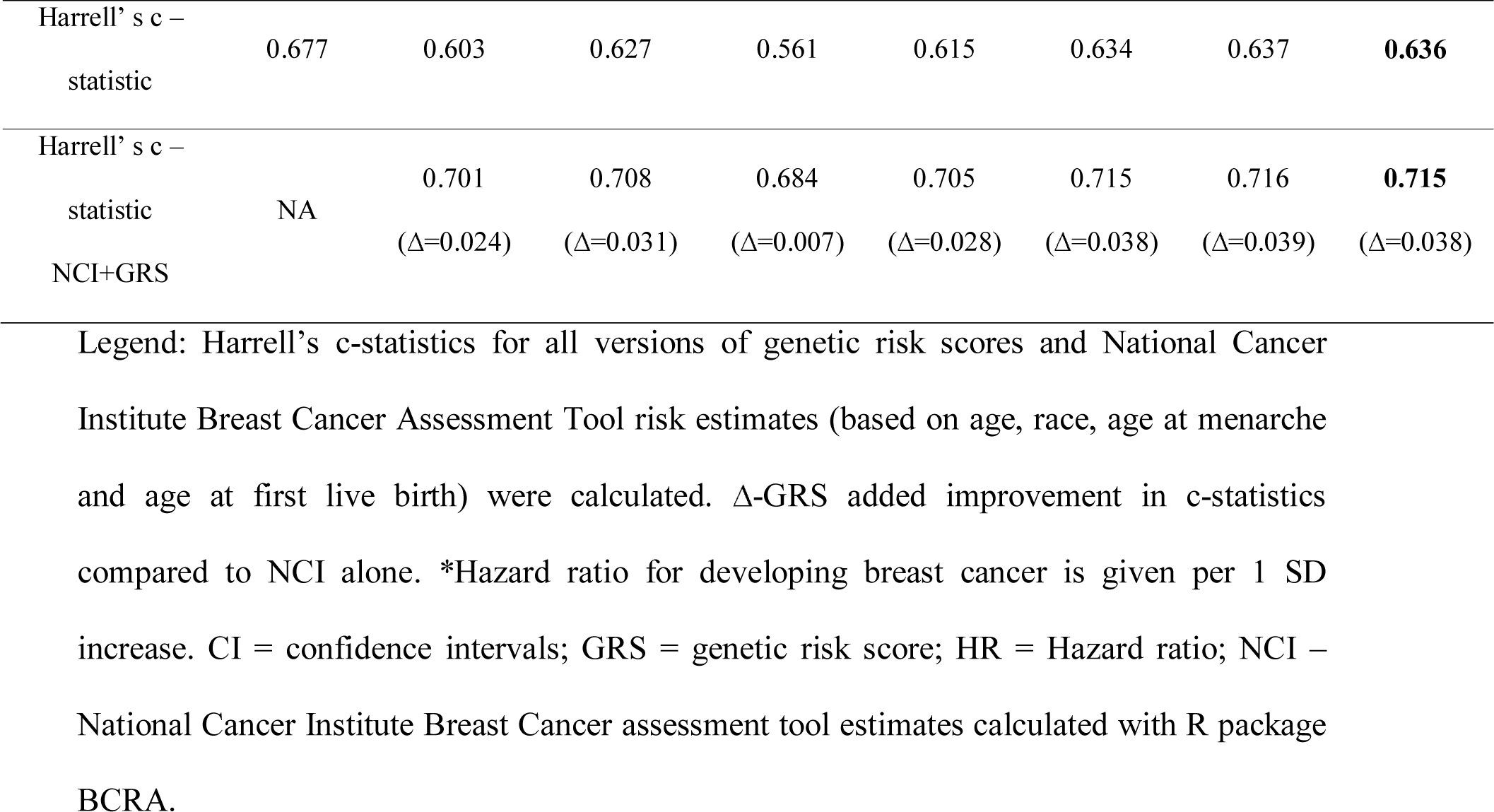
Analysis results for incident breast cancer in EstBB using different GRSs and metaGRSs.

### The score metaGRS_2_ and its potential for personalized breast cancer risk prediction

Women in the highest quartile of metaGRS2 distribution have 3.40 (95% CI 2.36 to 4.89) times higher hazard of developing BC than women in the lowest quartile. When the top quartile is further split into smaller percentiles (as seen on Figure 1), a strong risk gradient is seen also within this quartile. Namely, women in the top 5% of the metaGRS_2_ distribution have a Hazard Ratio (HR) of 4.79 (95% CI 3.02 to 7.58) for incident BC compared to women in the lowest quartile, whereas HR = 4.20 (95% CI 2.84 to 6.23) for women in the top 5% compared to all women with metaGRS2 below the median. When the highest 5% percentile is compared with the rest of the cohort (women below the 95^th^ percentile of metaGRS_2_), about three times higher hazard (HR = 2.73, 95% CI 1.92 to 3.90) is found.

**Figure 1.**
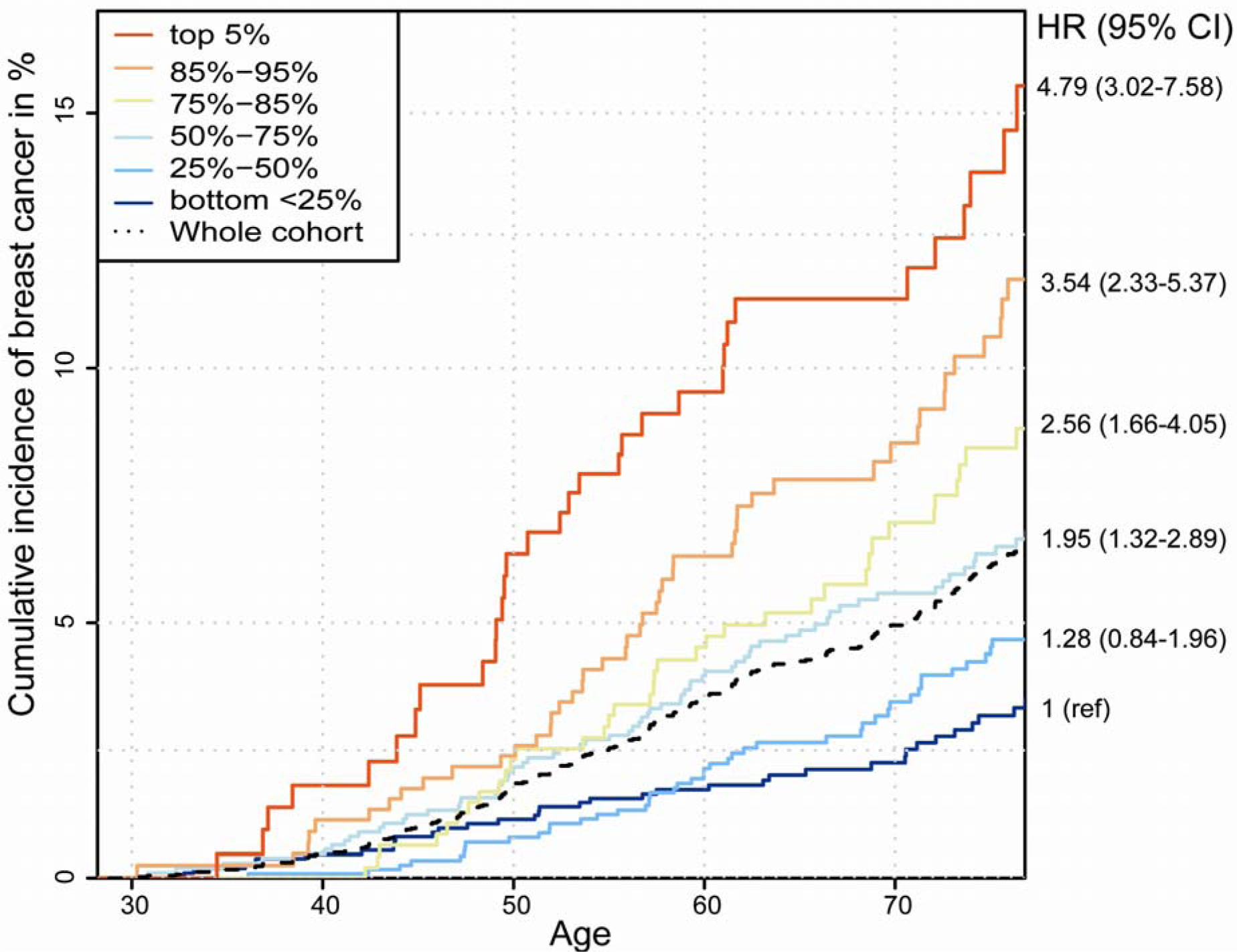
Cumulative incidence of BC in metaGRS_2_ categories among women within age 30-75 years. Legend: Cumulative incidence accounting for competing risks. Hazard ratios (HR) correspond to the comparison of several categories with the lowest quartile of metaGRS_2_.

As seen from Figure 1, the cumulative BC incidence by the age of 70 is estimated to be 12% (95% CI 7.7% to 16.3%) for women in the top 5% percentile of metaGRS2, 8.3% (95% CI 5.6% to 11.0%) for those between 85%-95% percentiles and 7.4% (95% CI 4.85% to 10.0%) for the women in 75%-85% percentiles. Cumulative BC incidence in the third, second and first quartile of the metaGRS2 distribution is estimated to be 5.8% (95% CI 4.4% to 7.3%), 3.6% (95% CI 2.4% to 4.8%) and 2.4% (95% CI 1.4% to 3.3%), respectively. No significant difference in BC hazard is seen between the two lowest quartiles (p = 0.26), with both of them having considerably lower incidence level than the cohort average (overall cumulative BC incidence estimated as 5.1% by the age of 70, 95% CI 4.5% to 5.8%).

### Correlation of GRSs

The correlations between seven scores varied between 0.3 to 1 (see Figure S2). While dividing individuals into 2 categories (“non-high” – GRS < 95th percent and “high” – GRS in top 5%) based on three GRSs (GRS_UK_, GRS_ONCO_ or GRS_75_), 87.7% (28547) of women were assigned to non-high category with all three scores. However, 12.4% (4010) of women belong to high category with at least one GRS. 0.33% (109) of women belonged to top 5% with all three scores compared to ∼10% (3240) of the women, who belonged into high category only with one score (Figure 2).

**Figure 2.**
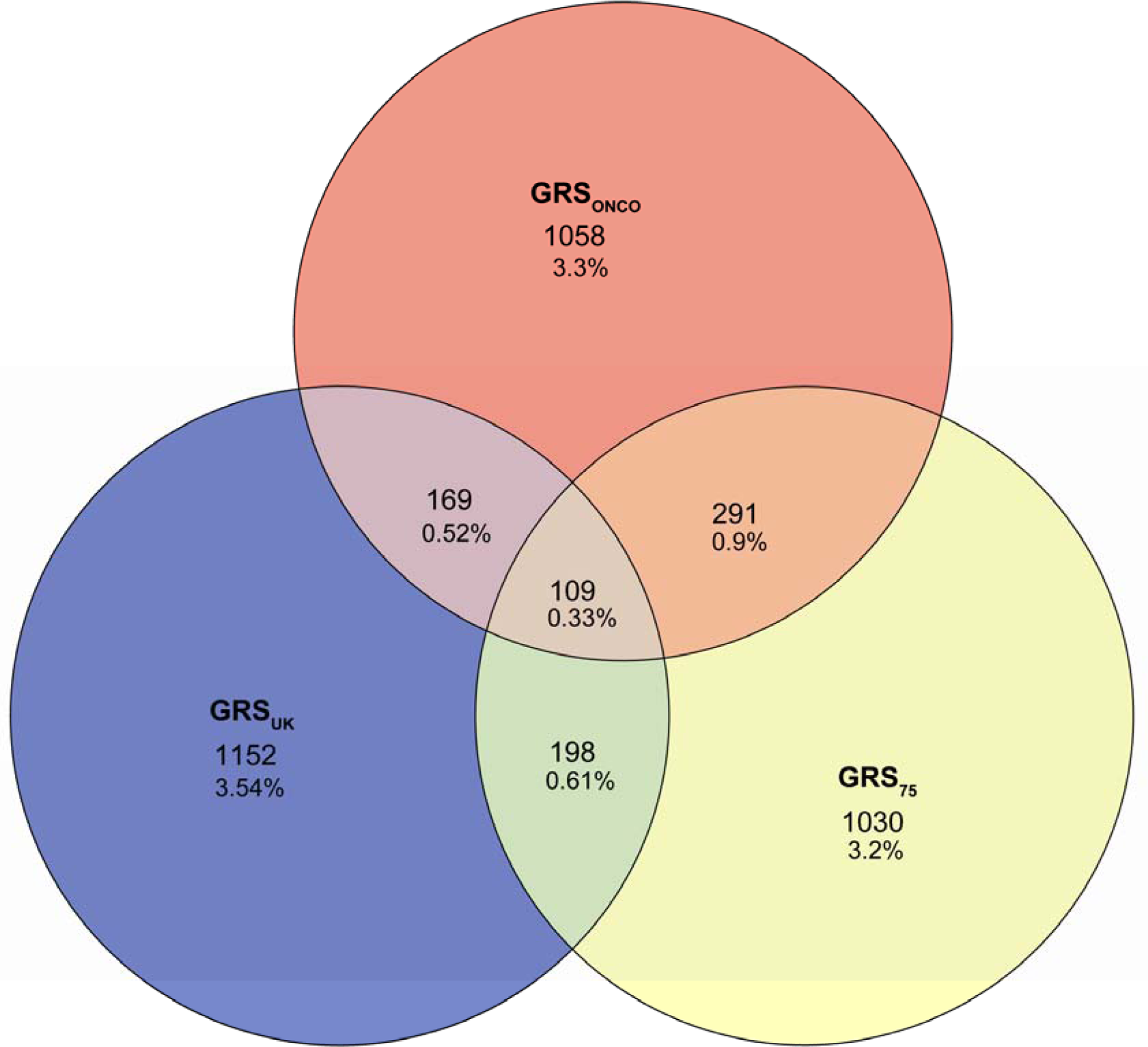
Division of Estonian Biobank women according to their genetic risk category. Legend: Women, who belong to top 5% at least with one out of the three genetic risk scores (GRSs: GRS_onco_, GRS_UK_, or GRS_75_), are represented on this graph. Number of women, who belong to top 5% only with one score, two scores or all three scores are given. Percentages are given per entire cohort.

### Associations of GRSs and other genetic and non-genetic predictors of breast cancer

Both family history as well as GRSs were strongly associated with BC status in UKBB, while the effects of GRSs were attenuated by less than 1% while adjusting for family history (Additional File 1, Table S2). Known BC risk factors were only weakly associated with in both UKBB and EstBB cohorts (Additional File 1, Table S3-S4). BMI and waist circumference were negatively associated with GRS_UK_ in both EstBB and UKBB, the association in EstBB was stronger for women under 50 years of age. Smoking status was positively associated with all GRSs except GRS_UK_ only in EstBB data. Age at menopause was associated with some GRSs in both cohorts but the effects were in opposite direction. No GRS showed association with any other type of cancer or overall mortality.

## Discussion

We demonstrate that a metaGRS that combines a multigenic and a polygenic GRS for breast cancer, metaGRS_2_, performs better than using either one of the previously published multigenic GRSs and also better than the best polygenic GRS alone. While in average about 5% of women in the EstBB cohort (as well as in the Estonian population) have been diagnosed with BC by the age of 70, women in the highest five percentiles of the metaGRS2 distribution have reached the same cumulative risk level (5%, 95% CI 2.1% to 7.8%) by the age of 49, thus more than 20 years earlier. It is also notable that women with metaGRS_2_ level below median reach such risk level (4.6%, 95% CI 3.6% to 5.6%) only by age of 79, thus almost 10 years later. This finding suggests that the polygenic risk estimate based on metaGRS_2_ could be an efficient tool for risk stratification in clinical practice, for targeted screening and prevention purposes.

Given that the potential benefits of non-selective BC screening within certain age categories (compared to potential harm from over diagnosis) are under serious discussion in medical community^23^, personalized approaches based on individual risk levels deserve further assessment. Ideally, those should integrate available information from clinical risk factors and also genetic information. The latter could include both moderate-and high-penetrance germline mutation testing, as well as polygenic risk scores. That approach is also supported by our findings, where considerable increase in c-statistics were observed while combining polygenic risk scores and NCI estimates together.

However, while incorporating a GRS in clinical BC prediction, one should keep in mind that a GRS represents a mixture of different pathways, but is still not likely to capture the heritable component completely. As our findings indicate that a GRS and family history have independent predictive effects on BC risk, accounting for individual’s genetic information and family history simultaneously in risk estimation could be recommended.

As depending on a GWAS that is used as a basis, different (and not necessarily highly correlated) GRSs can be produced, it is expected that those GRSs might emphasize the effects of different biological pathways. This hypothesis seems plausible in the light of several associations found between different GRSs and BC risk factors.

The fact that a metaGRS performs better than alternatives, suggests that the SNPs that are included in the multigenic GRS_75_ are potentially representing genetic pathways with stronger effect on the disease risk and the combined score will give them a stronger weight than the polygenic GRS alone. However, it also indicates that the SNPs included in the polygenic GRS_ONCO_ – but not in the GRS_75_ – have some predictive power and therefore one should not completely ignore them in an optimal GRS.

It remains an open question whether it is always the best practice to use metaGRS instead of several different genetic risk scores – if one can pinpoint biological mechanisms behind different scores, more optimal preventive strategies could be chosen. Still, until we are unable to convincingly link different GRSs with specific preventive measures, targeted prevention should be based on a GRS with the best possible overall predictive ability, such as the metaGRS_2_ proposed here.

One should also keep in mind that besides GRS there are genetic mutations such as BRCA1/2 known to be associated with very high familiar BC risk. Therefore, in practice, any genomic risk stratification should include search for high-risk genetic variants, or moderate risk variants, as well, if possible. In the high-risk mutation carriers, the clinical management could be based on the specific genetic (mendelian) variants, or if deemed useful in the future, a combination of mendelian variants and GRS levels, but it definitely needs further studies.

## Conclusions

In summary, our results show that an efficient polygenic risk estimate enables to identify strata with more than four-fold differences in BC incidence. This definitely calls for the development of personalized screening and prevention strategies that incorporate the GRS information, having the potential to considerably increase the benefits of nation-wide screening programs and reduce the existing controversies on their efficacy. However, one should be aware of the fact that a GRS is not uniquely defined – as more research accumulates, more efficient polygenic predictors could be developed that may re-categorize some previously stratified individuals into high or low risk groups. In addition, a GRS should ideally be combined with information on other genetic and non-genetic risk factors for best possible accuracy in risk assessment.

## List of abbreviations

BC: Breast Cancer
GWAS: Genome-Wide Association Study
GRS: Genetic Risk Score
EstBB: Estonian Biobank
UKBB: UK Biobank
SNP: Single Nucleotide Polymorphism
metaGRS: combination of several genetic risk scores, number in subscript indicates the number of original GRSs included
SD: Standard Deviation
HR: Hazard Ratio
OR: Odds Ratio
NCI: National Cancer Institute Breast Cancer
CI: Confidence Intervals

## Declarations

### Ethics approval and consent to participate

**EstBB**: All human research was approved by the Research Ethics Committee of the University of Tartu (approval 234/T-12), and conducted according to the Declaration of Helsinki. All participants provided written informed consent to participate in the Estonian Biobank.

**UKBB**: The UK Biobank study was approved by the North West Multi-Centre Research Ethics Committee (reference for UK Biobank is 16/NW/0274). All participants provided written informed consent to participate in the UK Biobank study.

### Consent for publication

Not applicable.

### Availability of data and material

We do not have ethical approval to share individual level genotype and phenotype data for Estonian Biobank. The data from UK Biobank were used under licence for the current study, and so are not publicly available. Researchers interested in Estonian Biobank can request the access here: https://www.geenivaramu.ee/en/access-biobank and access to UK Biobank can be requested here http://www.ukbiobank.ac.uk/resources/.

## Acknowledgements

This research has been conducted using the UK Biobank Resource under Application Number 17085.

## Additional files

In the file “Additional file 1” are four Supplementary Tables in *.xlsx format. Tables are labeled “S. Table 1-4”. The information included is following:

S. Table 1. Cohort characteristics of UK Biobank and Estonian Biobank.

S. Table 2. Associations of breast cancer and standardized GRSs in the UK Biobank (with and without adjustment of family history) and in Estonian Biobank without family history.

S. Table 3. Associations between GRSs and risk factors of breast cancer in Estonian Biobank.

S. Table 4. Associations between GRSs and risk factors of breast cancer in UK Biobank.

In the “Additional file 2” are Supplementary Figures and Methods in *.doc format. There are two supplementary Files and detailed information about genotyping, quality control, GWAS data management and statistical modelling for breast cancer risk factors and GRSs. The Supplementary figures are following:

Figure S1. Associations of GRSs with prevalent breast cancer in EstBB data.

Figure S2. Correlations between different genetic risk scores (GRSs).

